# CureSCi Metadata Catalog—finding and harmonizing studies for secondary analysis of hydroxyurea use for sickle cell disease

**DOI:** 10.1101/2024.08.15.608203

**Authors:** Xin Wu, Jeran Stratford, Karen Kesler, Cataia Ives, Tabitha Hendershot, Barbara Kroner, Ying Qin, Huaqin Pan

**Author notes:** Corresponding author: Huaqin Pan. These authors are shared first authors and contributed equally to this work (Xin Wu, Jeran Stratford).

## Abstract

**Objectives:** Sickle cell disease (SCD) is a rare group of inherited red blood cell disorders that affect hemoglobin, resulting in serious multi-system complications. The limited number of patients available to participate in research studies can inhibit investigating sophisticated relationships. Secondary analysis is a research method that involves using existing data to answer new research questions. Data harmonization enables secondary analysis by combining data across studies, especially helpful for rare disease research where individual studies may be small. The National Heart, Lung, and Blood Institute Cure Sickle Cell Initiative (CureSCi) Metadata Catalog is a web-based tool to identify SCD study datasets for conducting data harmonization and secondary analysis. We present a proof-of-concept secondary analysis to explore factors associated with discontinuation of hydroxyurea, a safe and effective first line SCD therapy, to illustrate the utility of the CureSCi Metadata Catalog to expedite and enable more robust SCD research.

**Methods:** We performed secondary analysis of SCD studies using a multi-step workflow: develop research questions, identify study datasets, identify variables of interest, harmonize variables, and establish an analysis method. A harmonized dataset consisting of eight predictor variables across five studies was created. Secondary analysis involved a generalized linear model was employed to identify factors that significantly impact hydroxyurea discontinuation.

**Results:** The CureSCi Metadata Catalog provided a platform to efficiently find relevant studies and design a harmonization strategy to prepare data for secondary analysis. Multivariate analysis of the harmonized identified that patients who are older, are female, had a history of blood transfusion therapy, had episodes of acute chest syndrome, and had the SC sickle cell genotype are more likely to stop hydroxyurea treatment.

**Conclusion:** This secondary analysis provides a template for how the CureSCi Metadata Catalog expedites dataset discovery of sickle cell studies for identifying relationships between variables or validating existing findings.

## Introduction

Sickle cell disease (SCD) is an inherited group of autosomal recessive disorders caused by mutations in both copies of the hemoglobin gene. Acute and chronic SCD-related complications often begin in infancy and manifest across multiple organ systems as hemolytic anemia, acute chest syndrome, stroke, eye problems, infections, and repeated episodes of pain called pain crises [1,2].

To date, there are few therapeutic options for people with SCD [2]. Hydroxyurea (HU) was the first disease-modifying therapy approved for use for SCD and continues as a first-line therapy for most individuals. Previous meta-analysis of nine randomized controlled trials evaluated HU use versus placebo, transfusion, or observation and found HU effective in decreasing the frequency of acute complications, preventing life-threatening neurological events, and improving survival rates for SCD patients [3–5]. However, there is insufficient evidence regarding the long-term risks and benefits and the effect of HU on and preventing chronic complications of SCD [5].

Several main barriers to initial HU use include safety and efficacy, side effects, fear of cancer, pregnancy and conception concerns, and decreased quality of life [6–13]. Factors associated with a patient’s stopping HU treatment are not well established. Preliminary analysis of additional factors associated with stopping HU can help elucidate the decision of HU use, which can help improve SCD patient treatment options and overall health. However, SCD is a rare disease [14], affecting fewer than 200,000 people in the United States and 20 million people worldwide [2,15]. As a result, the number of patients available to participate in research studies is often limited. This can lead to underpowered studies that may inhibit investigating more subtle or sophisticated relationships, including patient choices about initial and continued use of HU to manage SCD.

Data harmonization is an approach where similar independent studies are combined to create a larger study dataset for secondary and meta-analyses. This approach provides a more comprehensive understanding of the research topic, increased statistical power, and more reliable estimates of treatment effects. Harmonization of existing data sets can overcome challenges of heterogeneous populations and reduce the cost of research through reuse of existing data. Furthermore, data harmonization highlights gaps in knowledge to guide future research toward areas through secondary analysis. However, it is crucial to assess whether studies selected for secondary analysis are compatible in terms of patient populations, interventions implemented, and outcomes measured. Indeed, finding, preparing, and harmonizing study data for secondary analysis is one of the most time-consuming steps of the process.

The National Heart, Lung, and Blood Institute (NHLBI) Cure Sickle Cell Initiative (CureSCi) supported development of a Metadata Catalog (https://curesicklecell.rti.org/), a web-based open-access catalog of National Institutes of Health (NIH)-funded SCD studies used to identify datasets for reuse of existing data sets [16]. The Metadata Catalog’s browse and search capabilities can guide SCD researchers to discover relevant studies for pooled analysis by leveraging metadata at the study, patient Reported Outcomes (PRO) Measures, and data elements level. Associated datasets can then be requested through the NHLBI data repositories, BioLINCC or BioData Catalyst (https://biodatacatalyst.nhlbi.nih.gov/). Data harmonization enables investigators to address more subtle or sophisticated relationships by pooling datasets from multiple studies to overcome limitations imposed by small sample sizes.

In this study, we present a proof-of-concept data harmonization and secondary analysis to explore factors associated with discontinuation of HU treatment, using the CureSCi Metadata Catalog to identify datasets and streamline the process of conducting secondary analyses. We demonstrate this approach by investigating key factors that impact adherence to HU use. By leveraging the comprehensive and organized nature of a metadata catalog, we aimed to efficiently identify relevant studies and extract key information, prepare a harmonization strategy, and ultimately expedite our pooled analysis and enable a more robust exploration of factors associated with a patient’s stopping HU treatment.

## Methods

The workflow described in Figure 1 summarizes our steps for conducting data harmonization and secondary analysis of SCD study data. Although the arrows imply a certain order, the steps outlined here are more of a guideline and show the process by which our demonstration data harmonization and secondary analysis was conducted. For example, we list the research question as the first step, but the research question may evolve after surveying the types of datasets and variables available.

**Fig 1.**
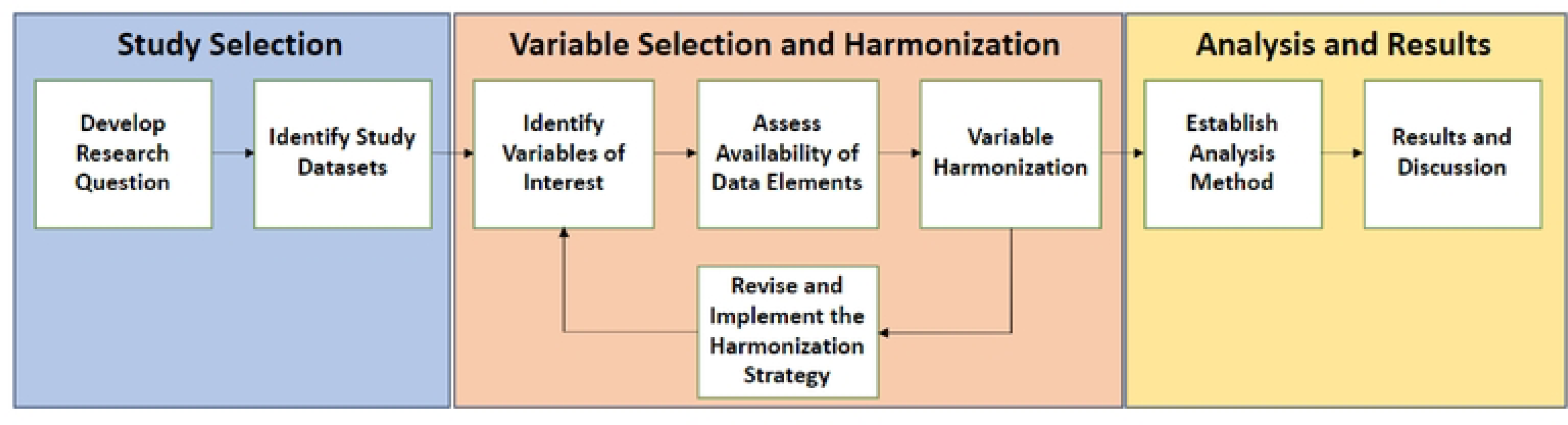
Process diagram of a data harmonization and secondary analysis workflow. A general step-by-step workflow for conducting data harmonization and secondary analysis can be applied to investigate key scientific questions.

### Developing a research question

Formulating a well-defined research question is critical for designing a focused and impactful research study. Several factors should be considered when defining the hypothesis, including addressing knowledge gaps within the existing literature, feasibility, availability of appropriate methodologies, and pressing clinical needs. A landscape analysis of SCD literature indicates that despite the substantial body of evidence demonstrating the benefits of HU, concerns remain that limit its use and adherence. While curating SCD studies for ingestion into the Metadata Catalog, we noted that many studies collected information regarding use and adherence to HU, as well as patient characteristics, symptoms, adverse events, and social determinants of health. Therefore, we hypothesized that data harmonization of existing studies and secondary analysis can produce a large-sample dataset where the factors that influence initiation of and adherence to HU can be determined.

### Study variables and harmonization strategy

#### Response variables—defining the HU user group

Various data elements can be collected for one concept, such as defining whether a patient is a current or past HU user, depending on how the data were collected in each study. A mapping schema that groups and identifies variables representing a concept should be specific to each dataset, with a goal of comparing and harmonizing across studies. Several sources of information were extracted from study protocols, data collection forms, data dictionaries, codebooks, and analysis plans hosted on the Metadata Catalog and used to develop the mapping schema, including (1) study inclusion and exclusion criteria, (2) treatment arm assignments, (3) participant responses on onboarding and longitudinal surveys regarding medication use, and (4) data on adverse events.

To facilitate cross-study comparisons, research participants were classified as current or past HU users. Current users are defined as patients who, at the time of data collection, began or continued an existing HU treatment plan. Past users are patients who were at one time on an HU treatment plan but chose to discontinue taking HU.

#### Predictor variables

Adherence to therapeutic regimens is influenced by a complex interplay of factors, including tolerability, methods of treatment administration, and adverse events. Furthermore, individual characteristics, including age, sex, race, ethnicity, and genetic polymorphisms, may influence treatment response and thus contribute to overall treatment effectiveness and adherence to the therapy. In addition to outcomes, it is important to control for factors that may be related to, or confound, the outcome of interest or may be different across study populations. Several factors have been reported in the literature to be associated with a patient’s decision to start or continue taking HU. For instance, HU studies focused on pediatric SCD populations demonstrate the association with age [17,18], as well as gender-specific effects such as impairing seminal fluid concentration and diminishing ovarian reserves [19]. Furthermore, HU has been shown to be associated with adverse pregnancy outcomes [13]. In addition, HU use may be influenced by SCD severity and assessed using indicators for stroke, acute chest syndrome, pain severity, and SCD genotype [1,5,7,8]. Finally, effective use of alternate therapies (e.g., blood transfusion therapy, pain medication) is known to reduce HU use.

### Identifying study datasets

The CureSCi Metadata Catalog offers researchers the ability to explore 27 SCD studies, including study protocols, manuals of operation, study summaries, case report forms, data dictionaries, codebooks, and location of full datasets. Users can browse and search curated study variables and PRO measures using the “Browse Studies by Data Element” and “Browse Studies by PRO Measure” tools, respectively. These tools allow users to hierarchically browse data elements and PRO measures that have been curated into descriptive domains and subdomains.

The Participant Frequency Tool (https://curesicklecell.rti.org/FreqTool) allows users to assess the extent of missing data across studies and selected variables. The resulting output is a count of participants who have data for the selected variables, which can be used to assess feasibility of further analyses based on previously performed power calculations.

### Variable harmonization

Before combining data across multiple studies for analysis, the study variables must first be harmonized. Data harmonization is the process of taking data from various sources and making them consistent, often requiring data transformation to ensure that measures are reported in consistent units and a unified data format to ensure accurate data interpretation. This nontrivial process requires intimate knowledge of the data and the collection mechanism used, often requiring abstraction to standardize the meaning and representation of data from different sources. Identification of common data elements (CDEs) greatly accelerates this process.

To ensure data comparability across studies included in this data harmonization analysis, we conducted a comprehensive variable extraction process. All relevant variables from each study for each response or predictor variable were identified. Subsequently, we determined the most frequently employed collection approach or coding scheme for each variable. This most common approach was then adopted as the standard for harmonizing the remaining studies. Study variables were compared with the chosen standard and categorized as identical, comparable, or unrelated. Identical variables used the collection standard or very similar study descriptions and answer responses requiring little to no further harmonization. Comparable variables required some modification of the answer scales or response options to be harmonized to the standard. Unrelated variables had answer scales that could not be mapped or transformed to the standard. For instance, variables reporting pain for different time scales (chronic vs. acute) and body parts (head vs. joints) could not be combined. “Unknown” and “not applicable” responses were dropped from the analysis.

### Data extraction and harmonization

For each study, the Metadata Catalog provides the location of the data and the mechanism for requesting access to the data. Links provided in the Metadata Catalog to datasets deposited in repositories (e.g., BioLINCC) were used to download study datasets after submission and acceptance of the data access agreement. Additional datasets were procured through data access requests directly to the study investigators, who are listed with their affiliated institute in the Metadata Catalog. SAS 9.4 was used to extract study variables identified in the harmonization strategy from the data files. We first verified that the question and responses for each variable matched what was reported on the data collection form. Data transformation instructions outlined in the harmonization strategy were then implemented. After harmonization, study data were combined into a single dataset. Variable response rates were tabulated to identify any missing data and to validate that each response level had sufficient counts to support the statistical model. Finally, the dataset was exported into a single comma-separated values file for additional processing and analysis.

### Statistical analysis

Logistic regression was used to investigate the factors that influence adherence to an HU treatment plan. The HU user group (current vs. past HU users) was modeled as a function of the predictor variables. First, we performed a univariate analysis where HU use was modeled as a function of each individual predictor, then a multivariable model controlling for all predictors was modeled, then was refined using a backward elimination approach.

Backward selection was performed in SAS 9.4 to produce a parsimonious model. The log likelihood test was used to determine appropriate model fit. HU use was modeled as a factor of all eight predictors (S1 and S4 Tables).

## Results

### Research question

HU is considered a safe and effective therapy for SCD. Although the effect of HU use for SCD is well studied [3–5], an in-depth investigation of factors associated with discontinuation of HU has not been conducted. For this study, we sought to investigate the hypothesis that factors such as age, gender, race, ethnicity, SCD genotype, acute pain episodes, pain severity, stroke, acute chest syndrome, pain medication use, and transfusion history influence continued use of HU to manage SCD.

### Study selection

After defining the research question, we used the CureSCi Metadata Catalog to identify potential studies for the data harmonization analysis. To identify relevant variables for our data harmonization analysis, we used the catalog’s “Browse Studies by Data Element” feature. Searching in the “Therapeutics” domain and the “Medication” subdomain, we identified 195 variables across 20 studies specifically related to HU. Of the remaining 20 studies, we had participant data for 8 that we could use for immediate data analysis. We performed additional study selection by querying “hydroxyurea history” and “hydroxyurea current use.” As a result, we selected the eligible studies that had participant data and their appropriate data collection phase and eliminated 2 studies that lacked key variables and 1 transplant trial because effects were mainly due to transplant rather than HU (Table 1). This reduced the number of studies to 5, allowing us to narrow down the list of data sources for the secondary analysis (Table 1).

**Table 1.**
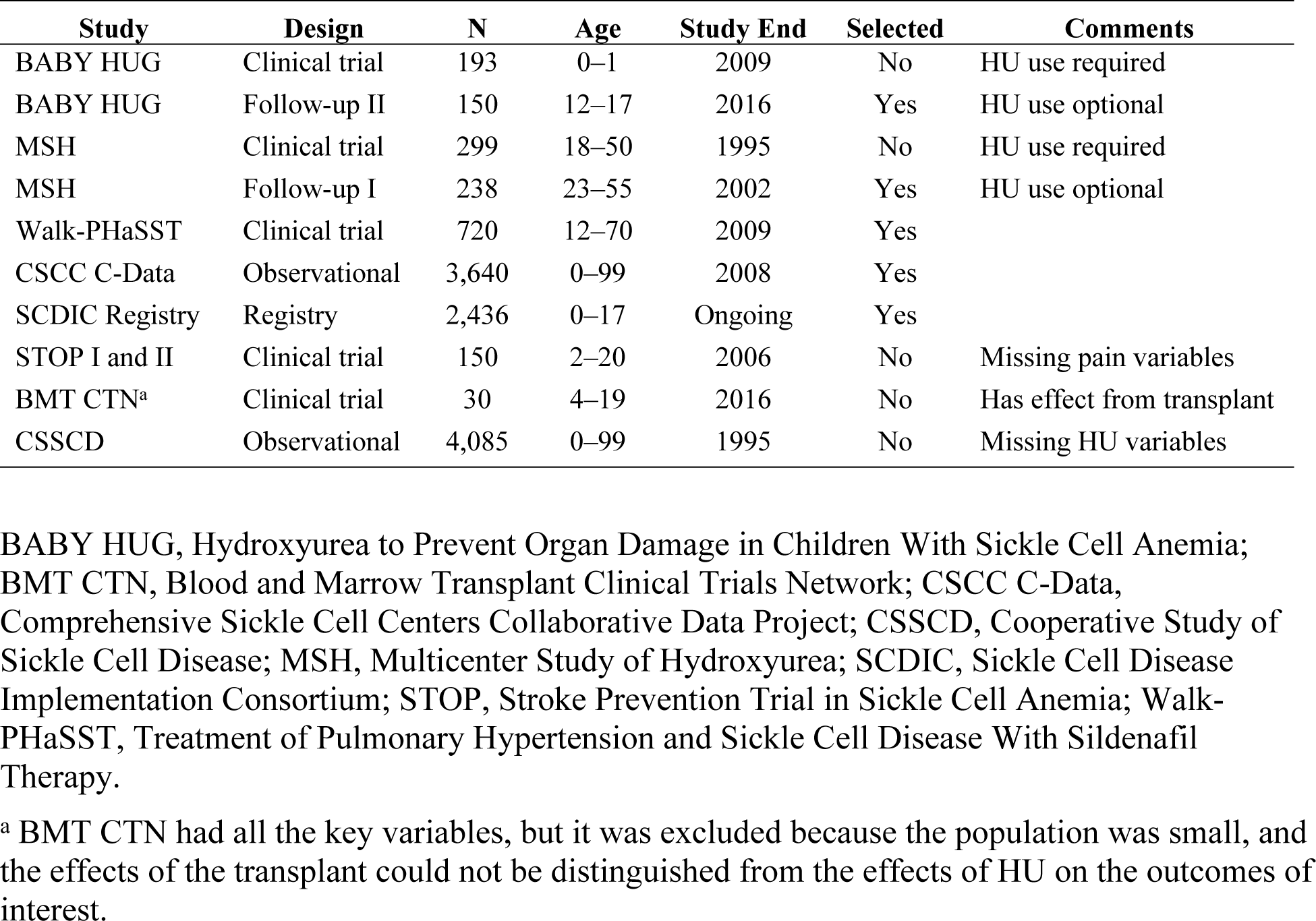
Selecting study datasets for the HU use harmonization and secondary analysis using the CureSCi Metadata Catalog.

Final selection of studies to include in the secondary analysis was determined by identifying which of the filtered studies that collected data on HU also collected data on each of the proposed factors (Table 1).

Two of the selected studies (BABY HUG and Multicenter Study of Hydroxyurea [MSH]) were multiphase studies consisting of a main clinical trial and observational follow-up periods. We identified variables collected at each phase using the Metadata Catalog. Because not all datasets included in the secondary analysis provided longitudinal data, we employed a specific selection strategy to identify a single phase from the longitudinal cohorts to provide a representative snapshot. This strategy involved choosing the collection phase of these studies where the response variable and all predictor variables were measured. This ensured that the analyzed predictors would correspond temporally to the measured responses, minimizing potential biases introduced by lagged relationships between variables. For BABY HUG, the initial clinical trial centered on the effects of HU in infants, meaning patients were in the drug group or the placebo group and did not have a choice between selecting their study group and discontinuing treatment. We chose the second follow-up period as the source of our data because patients were given the choice to continue HU therapy in this phase (these data were not collected in the Phase 1 follow-up) and were now old enough to give more accurate pain assessments. For MSH, like BABY HUG, patients enrolled in the clinical trial were in the HU drug group or the placebo group. Therefore, we chose to use data from follow-up I, where patients were given a choice to stop or continue HU use.

### Variable selection and harmonization

After compiling the initial list of predictors and covariates in the harmonization design, we documented the presence of each variable in each eligible study (Table 2). We then reviewed variable descriptions and answer values, determined whether they could be harmonized for the secondary analysis, and assessed variable availability in each study. As a result, we selected the following variables for the harmonization and secondary analysis (S1 Table) and eliminated three variables (insurance, marital status, and seizure) because of inconsistent or absent data collection across studies (Table 2).

**Table 2.**
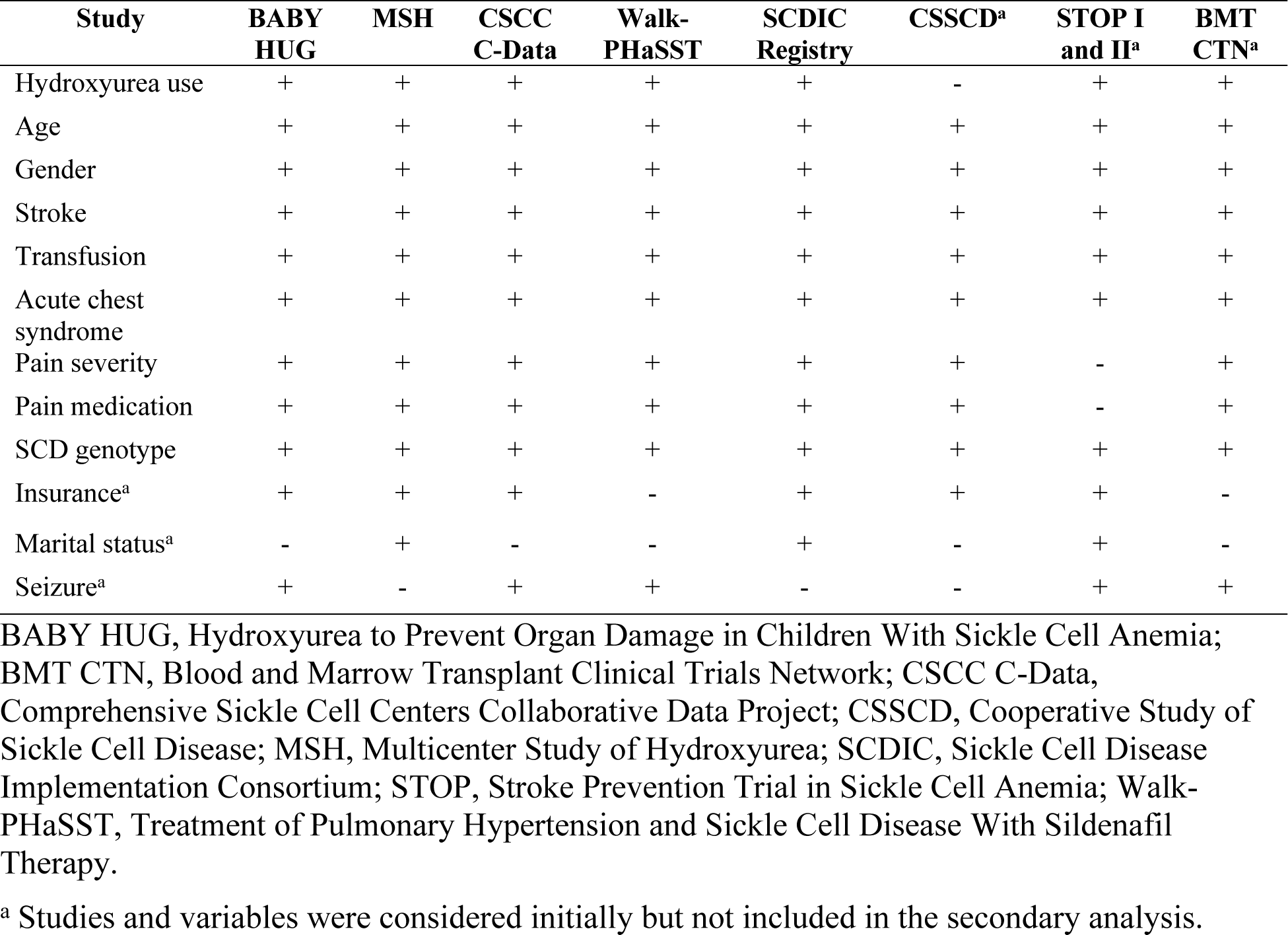
Eligible studies from the CureSCi Metadata Catalog and their available predictor variables.

Data harmonization was performed individually for each study dataset and included standardizing measurement scales and recoding variable responses. This standardization process minimizes potential biases introduced by methodological heterogeneity and facilitates a more robust analysis. Harmonization efforts for secondary analysis can vary considerably, depending on the nature of the data discrepancies. Simple adjustments, like rescaling scores from a 0 to 10 to a 0 to 5 range, are relatively easy and reasonable to achieve. However, harmonization becomes increasingly challenging for variables whose collection had more substantial methodological differences. This includes studies that ask entirely different questions, have significantly varied follow-up times, or use fundamentally different measurement tools. For example, every study that collects pain reports it differently. To harmonize these variables, we first checked the pain context (general pain vs. specific body part or time frame) and that the responses were of the same type (numerical scale rather than open ended). Furthermore, numerical measurement of pain severity needed to be harmonized to the same scale. By using the CureSCi Metadata Catalog, we were easily able to find all pain variables for each study and determine whether they fit our criteria and were harmonizable. Variables are organized into domains, and users can simply hover over the variable name to show the question associated with the variable and all the response options. This feature enabled us to efficiently filter pain variables in each study that asked about recent general pain attacks on a numerical scale. A summary of the harmonized variables can be found in Table 3, and detailed descriptions of the harmonization strategy and the participant distribution for each variable across studies can be found in S1 and S2 Tables, respectively. A detailed example of how HU users were harmonized can be found in S1 Fig.

**Table 3.**
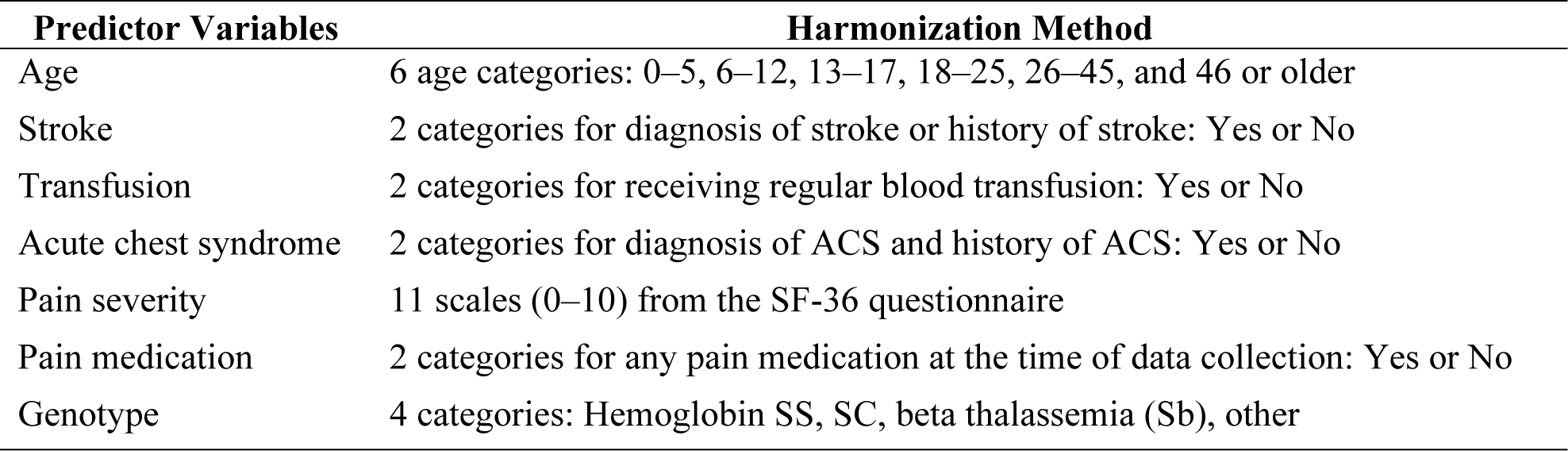
Predictor variables used in analysis and the methods used to harmonize to the categorical variables.

### Requesting study data

After we identified relevant studies, the next step was to complete the data access request. Links to submit applications for data access were collected from the Catalog’s “Browse Studies” feature under “Data Location.” Three of the studies (MSH, Walk-PHaSST, and BABY HUG) were available for request through BioLINCC; the Sickle Cell Disease Implementation Consortium (SCDIC) and the Comprehensive Sickle Cell Centers Collaborative Data Project required emailing the study principal investigator. Having these resources readily available in the Metadata Catalog resulted in an expedited submission process.

### Factors associated with discontinuation of HU treatment

Once the data had been obtained, the previously developed harmonization strategy needed to be implemented for each individual study.

Based on the type of data and variables used, an appropriate analytical pipeline should be established to analyze the harmonized data. For our HU analysis, the data were mainly categorical, which is why we opted for a generalized linear model (GLM; see the Methods section).

As part of the secondary analysis workflow (Fig 1), we performed a univariate GLM for each of the predictors and a multivariate GLM using all eight predictors. Identification of the most significant predictors in the multivariate GLM was performed by applying backward elimination (see the Methods section). Five of the eight predictor variables yielded statistically significant results (Fig 2 and Table 4)—age, gender, blood transfusion, stroke, and genotype. These findings are summarized in S3 and S4 Tables.

**Fig 2.**
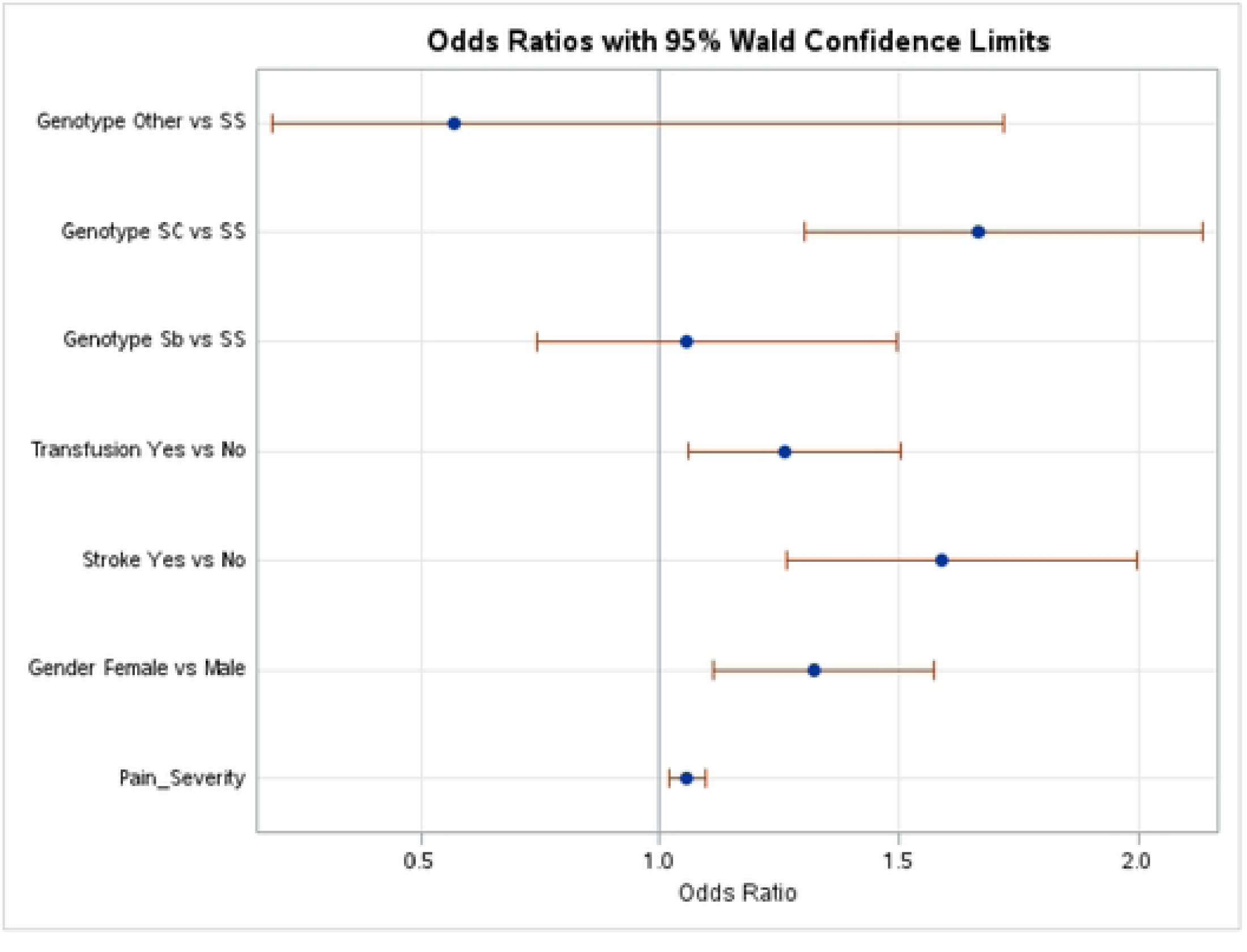
Odds ratios of factors that influence adherence to HU therapy for SCD patients. Forest plot showing the odds ratios (ORs) and 95% confidence intervals (CIs) for the association between hemoglobin genotype, blood transfusion history, stroke, gender, and pain severity with discontinuing HU therapy from a multivariate logistic regression analysis. Circles represent variable ORs, with lines indicating CIs.

**Table 4.** Results of the final model with five significant predictors for discontinuing HU.

| Characteristic | Level | Current HU User | Past HU User | Odds Ratio (95% CI) | P Value |
| --- | --- | --- | --- | --- | --- |
| Genotype | SS | 1,966 | 746 | (Reference) |  |
|  | Sb | 187 | 66 | 1.054 (0.742, 1.496) | 0.7841 |
|  | SC | 291 | 172 | 1.666 (1.300, 2.135) | 0.0028 |
|  | Other<sup>a</sup> | 29 | 7 | 0.569 (0.188, 1.718) | 0.1841 |
| Transfusion | No | 1,186 | 481 | (Reference) |  |
|  | Yes | 1,247 | 499 | 1.262 (1.058, 1.505) | 0.0096 |
| Stroke | No | 2,062 | 378 | (Reference) |  |
|  | Yes | 746 | 230 | 1.590 (1.264, 1.999) | <0.0001 |
| Gender | Male | 1,256 | 421 | (Reference) |  |
|  | Female | 1,219 | 571 | 1.323 (1.112, 1.574) | 0.0016 |
| Pain severity |  | Continuous variable |  | 1.055 (1.017, 1.094) | 0.0043 |

Summary statistics include the odds ratio, 95% confidence interval, and *p* value. The intercept represents the odds ratio for discontinuing HU for the reference group (males with an SS genotype who have not had transfusions or stroke and who reported a pain severity index of 0).
<sup>a</sup> SD, SO, SE, or other unspecified genotypes
<sup>b</sup> CI - Confidence Interval

Female SCD patients were more likely to discontinue HU therapy (odds ratio [OR] = 1.323), consistent with what has been reported in the literature with regard to the desire for pregnancy. An SC genotype had the largest OR among the significant predictors (OR = 1.666), indicating that individuals with an SC genotype were more likely to have discontinued use of HU compared with individuals with an SS phenotype. There was no association with Sb or “other” genotypes and discontinuing HU use. Similarly, individuals with a history of receiving transfusions were also more likely to be past users of HU (OR = 1.26). Individuals with severe pain or a history of stroke had increased odds of discontinuing HU (ORs = 1.055 and 1.590, respectively). The remaining predictors (acute chest syndrome, pain medication, and categorical age) were removed in the backward elimination process, with no statistically significant associations with HU use.

## Discussion

Decades of SCD research have yielded a rich corpus of data that serves as a valuable resource that can be leveraged to inform and conduct future research. Because SCD is a rare disease, secondary analysis of existing data helps overcome small sample size limitations, enable exploration of new hypotheses, identify previously unseen trends, uncover the underlying biology, and determine new therapeutic strategies. This data-driven approach can significantly accelerate progress in understanding and ultimately treating SCD.

In this work, we provide a flexible framework for conducting harmonization and secondary analysis of SCD studies. A curated metadata catalog is an asset when designing and planning a pooled analysis. The CureSCi Metadata Catalog serves as a specialized search engine for SCD research data, significantly streamlining the often arduous process of accessing relevant datasets. The Metadata Catalog expedites this process by offering a centralized platform for searching and filtering studies based on predefined data elements and PRO measures relevant to secondary and meta-analyses, including demographics, clinical history and physical examinations, environmental exposure, health care utilization, medical imaging and tests, therapeutics, and quality-of-life measures.

Traditionally, researchers might obtain study data, only to discover that the variables they require are not actually present. The Metadata Catalog also enables researchers to pinpoint studies containing relevant study variables before investing significant time and resources in data access agreements. Using the Metadata Catalog reduces this risk by providing curated lists of collected study variables. This transparency also allows researchers to efficiently design harmonization strategies while the data access request is being reviewed. In addition, the Metadata Catalog has a “Participant Frequency Tool” for researchers to quickly identify study sample sizes and response distributions for several curated key variables such as age, gender, ethnicity, pain, and more. Thus, an appropriate abstraction definition and data transformations required across studies can be coded and available for implementation shortly after the data are received.

Harmonizing data elements collected from different studies is an essential first step in performing a secondary or meta-analyses. Use of CDEs and PRO measures would enable pooled analysis without harmonization. In this study, we found that use of CDEs and PRO measures is lacking in many studies, especially those conducted more than 10 years ago. A success story is the use of PRO measures for “pain severity,” where a general 36-item Short Form Health Survey was used in four studies, and an SCD-specific Adult Sickle Cell Quality of Life Measurement Information System Pain Episode was used in one study (SCDIC) [20]. To harmonize these two PRO measures, we converted a 6-scale measure to an 11-scale measure. Some important factors such as emotional and educational impact were excluded from the final analysis because of the lack of CDEs. In other cases, we had to use some assumptions to harmonize the data elements. For example, we harmonized current and past blood transfusions into the transfusion predictor variable, along with all transfusion amounts greater than 0. Similarly, we harmonized all stroke incidents regardless of type, location, or severity into the stroke predictor. Based on this analysis, the adoption of CDEs and PRO measures is imperative for new and ongoing studies being designed or considered. This will minimize the effort required for harmonization for secondary use of the study data and maximize the study impact by making the results of the study comparable with those of other studies. CDEs will enable researchers to take advantage of existing resources to perform reproducibility studies, answer new scientific questions, and identify areas for future research. This is a best practice to comply with the Findable, Accessible, Interoperable, and Reusable (FAIR) principles and the NIH Policy for Data Management and Sharing. To this point, the NHLBI CureSCi has developed two data standards resources: (1) a Case Report Form Library and CDE Catalog (https://curesickle.org/datatools-overview) for prospective SCD studies, and (2) a tool annotating PRO measures for retrospective studies at “Browse Studies by PRO Measure” (https://curesicklecell.rti.org/MetaCatalog) [16]. The NHLBI SCDIC Registry (Phase II) study has adopted 21 PRO measures, as summarized in the “Report by PRO Measures” tool (https://curesicklecell.rti.org/Report/DetailPROMeasureReport). The detailed PRO measures adopted by the studies in the CureSCi Metadata Catalog can be browsed in the “Browse Studies by PRO Measure” tool (https://curesicklecell.rti.org/MetaCatalog).

To demonstrate the effectiveness of the harmonization and secondary analysis framework, we investigated factors affecting a patient’s decision to stop HU treatment. By combining patient data from five studies, we performed a GLM using 7,288 samples. We discovered five statistically significant factors associated with stopping HU treatment—gender, blood transfusions, stroke, pain severity, and SC genotype (Table 4). Our findings suggest that older patients, female patients, patients on current or past blood transfusion treatment, patients who have experienced acute chest syndrome, and patients diagnosed with the SC genotype all are associated with being more likely to stop their HU treatment. Patients may begin to suffer from more severe effects of SCD with increased age, which may prompt them to transition to more drastic treatment plans [21]. Similarly, acute chest syndrome is a severe symptom of SCD, similarly patients with the SC genotype present with more severe complications compared with the more common SS genotype, indicating that patients in both categories may also transition to more drastic treatments [5]. Female patients, on the other hand, may stop HU treatment due to its potential effects on fertility [2,5]. Finally, blood transfusion is a mutually exclusive treatment plan with HU and therefore an obvious factor in stopping HU use [22].

This study used available datasets to demonstrate the feasibility of conducting meta-analyses. The analysis yielded five factors significantly associated with HU use; these are not causal relationships. The analysis model reported here also lacks consideration for complex interactions and inclusion of other potentially important factors. For example, quality of life is an important underlying factor when studying health outcomes, although it was not included in our analysis because of limited data and was not harmonizable across studies.

In conclusion, the CureSCi Metadata Catalog expedites dataset discovery that enables pooled analysis of SCD studies for discovering more subtle or complex relationships or validating existing findings.

## Funding

This research was funded by the National Institutes of Health (NIH) Agreement OT3 HL147798. The views and conclusions contained in this document are those of the authors and should not be interpreted as representing the official policies, either expressed or implied, of the NIH.

## Data Availability

Data used for this study are available at BioLINCC.

## Notes

### Competing Interest Statement

The authors have declared no competing interest.

